# Patient-derived xenograft culture-transplant system for investigation of human breast cancer metastasis

**DOI:** 10.1101/2020.06.25.172056

**Authors:** Dennis Ma, Grace A. Hernandez, Austin E.Y.T. Lefebvre, Hamad Alshetaiwi, Kerrigan Blake, Kushal R. Dave, Maha Rauf, Justice W. Williams, Ryan T. Davis, Katrina T. Evans, Madona Y.G. Masoud, Regis Lee, Robert A. Edwards, Michelle A. Digman, Kai Kessenbrock, Devon A. Lawson

## Abstract

Metastasis is a fatal disease where research progress has been hindered by a lack of authentic experimental models. Here, we develop a 3D tumor sphere culture-transplant system that facilitates the expansion and engineering of patient-derived xenograft (PDX) tumor cells for functional metastasis assays *in vivo*. Orthotopic transplantation and RNA sequencing analyses show that PDX tumor spheres maintain tumorigenic potential, and the molecular marker and global transcriptome signatures of native tumor cells. Tumor spheres display robust capacity for lentiviral engineering and dissemination in spontaneous and experimental metastasis assays *in vivo*. Inhibition of pathways previously reported to attenuate metastasis also inhibit metastasis after sphere culture, validating our approach for authentic investigations of metastasis. Finally, we demonstrate a new role for the metabolic enzyme NME1 in promoting breast cancer metastasis, providing proof-of-principle that our culture-transplant system can be used for authentic propagation and engineering of patient tumor cells for functional studies of metastasis.

## Introduction

Metastasis is the cause of >95% of breast cancer patient mortality, causing death in >40,000 women per year^1,2^. There are limited effective treatments for metastasis, and the identification of new therapeutic targets has been hindered by a lack of experimental models that faithfully recapitulate metastatic disease. Most metastasis research has been conducted with a handful of genetically engineered mouse models (GEMM) and cell lines that have limitations for accurately modeling metastasis in humans^3,4^. PDX models, where patient tumors are propagated in mice, offer increased authenticity as they maintain intratumoral heterogeneity, molecular markers and pathological characteristics of the original patient tumor^5–7^. PDX models have been used extensively in pre-clinical studies for drug testing *in vivo*, but their use in metastasis research has been limited by technical challenges. Functional studies of metastasis often involve genetic or pharmacologic perturbation of cancer cells in culture followed by transplantation *in vivo* to determine the effect of a gene or pathway of interest on metastatic dissemination. Although studies have reported transient cultures of PDX cells for drug testing *in vitro*^8–10^, they are typically only maintained for 24-72 hours and not used for viral engineering or transplantation. Development of a robust method for the propagation, engineering and transplantation of PDX tumor cells would enable new functional investigations of metastasis using patient tumor cells.

Although human cancer cell lines are typically maintained in 2D culture, previous work has shown that 3D cultures provide several advantages for cultivating primary tumor cells. Since they are 3D, they more closely recapitulate the physiologic conditions found in normal tissues where cell phenotype and function is heavily influenced by cell-cell and cell-extracellular matrix (ECM) interactions^11^. 3D conditions can be generated using various conditions, such as the hanging drop method used to maintain embryonic stem (ES) cells^12–14^, suspension in low-adhesion conditions as used for neurosphere cultures^15^, or embedding in ECM scaffolds such as collagens, Matrigel, or synthetic hydrogels, which have been used to enrich for cancer stem cells (CSCs)^16^. Here, we screened several 3D culture conditions and present an optimized method for propagation of PDX cells as tumor spheres. RNA sequencing (RNA-seq) analysis shows PDX sphere cells maintain gene expression signatures characteristic of uncultured PDX tumor cells. Sphere cells can be engineered with lentivirus and produce tumors that recapitulate native PDX tumors following orthotopic transplantation *in vivo*. Importantly, we find that PDX sphere cells yield robust spontaneous metastasis from orthotopic tumors, as well as experimental metastasis following intravenous (i.v.) and intracardiac (i.c.) delivery *in vivo*. Inhibition of known metastasis-promoting pathways attenuates metastasis following tumor sphere culture, validating our approach for authentic investigations of metastasis from patient tumor cells. Finally, we investigate the role of nucleoside diphosphate kinase A (NME1), a gene we previously found upregulated during metastatic seeding of PDX tumor cells^17^. We find that NME1 overexpression promotes metastasis of orthotopically-transplanted, cultured PDX cells, providing proof-of-principal for the value of this culture-transplant system for facilitating functional analyses of new genes of interest in patient tumor cells.

## Results

### Development of a 3D culture system for viable propagation of PDX cells as tumor spheres *in vitro*

We screened several culture conditions to develop an optimal method for viable expansion of PDX tumor cells *in vitro* **(Fig. 1a)**. We assessed two 3D growth methods: 1) suspension, where cells are grown in ultra-low attachment (ULA) plates^18,19^, and 2) Matrigel, where cells are grown in a basement membrane-rich semi-solid substratum^20^ **(Fig. 1a)**. We also tested two media conditions: 1) Mammary Epithelial Cell Growth Medium (MEGM), which supports short-term cultures of PDX cells for drug testing^8^, and 2) EpiCult™-B (EpiCult), which is commonly used for breast epithelial cell culture^21,22^ **(Fig. 1a)**.

**Figure 1.**
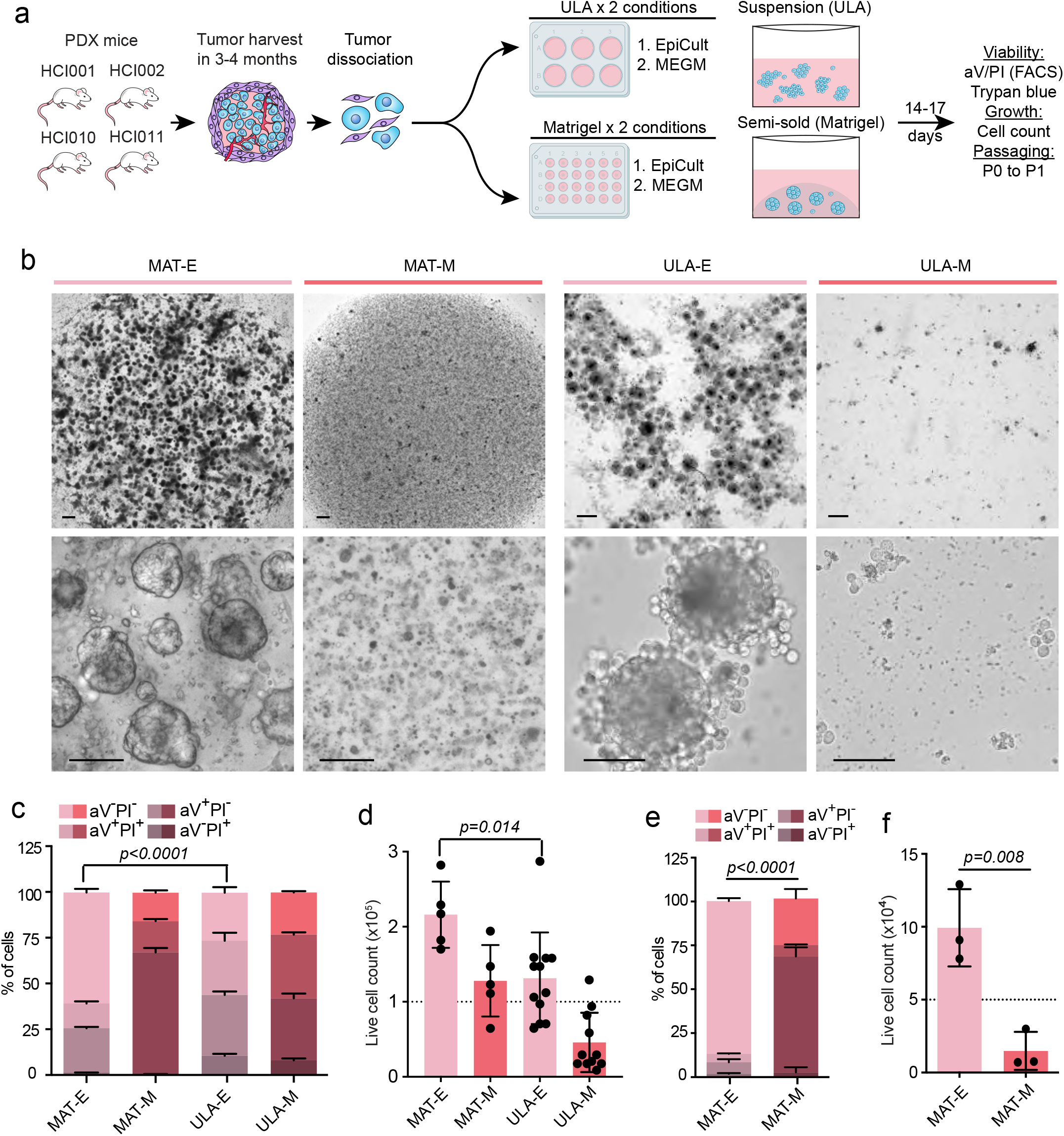
Comparison of 3D culture methods for viable propagation of PDX tumor cells *in vitro*. **(a)** Schematic summarizes culture conditions compared and readouts for PDX cell viability, growth and passaging capacity. **(b)** Representative brightfield images show spheroid structures generated 7-14 days after plating 2.5 × 10^5^ HCI010 cells in MAT-E, MAT-M, ULA-E, and ULA-M conditions. Scale bars = 300 µm (top row), 100 µm (bottom row). **(c)** Flow cytometry analysis of HCI010 cell viability post culture by aV and PI staining. Bar graph shows percent of aV-PI-(live), early apoptotic (aV+PI-) and dead (aV^+^PI^+^, aV-PI^+^) HCI010 cells in each condition. *P*-value determined by unpaired t-test comparing frequency of aV-PI-viable cells in MAT-E vs ULA-E, n=3 per condition. Data represented as mean ± s.d. **(d)** Bar graph shows total viable HCI010 cell number in each condition at the conclusion of culture. Cells were plated at 1.0 × 10^5^ cells/well (dashed line), and quantified nine days later by trypan blue exclusion. *P*-value determined by unpaired t-test comparing number of live cells in MAT-E vs ULA-E, n=5-11 wells per condition. Data represented as mean ± s.d. **(e)** Flow cytometry analysis of HCI010 cell viability in each condition post passaging by aV and PI staining. HCI010 cells were cultured for 2 weeks, dissociated, re-plated at 5.0 × 10^4^ cells/well in MAT-E and MAT-M and analyzed 21 days later. *P*-value determined by unpaired t-test comparing aV-PI-viable cells, n=3. Data represented as mean ± s.d. **(f)** Bar graph shows total viable HCI010 cell number in each condition after passaging as described in (e). Cell number was determined by trypan blue exclusion, and dashed line indicates initial cell number plated. *P*-value determined by unpaired t-test, n=3. Data represented as mean ± s.d.

We used the previously established breast cancer PDX models, HCI001, HCI002, HCI010 (ER-PR-Her2-; basal-like), and HCI011 (ER^+^PR^+^Her2^−^, luminal B)^5^ **(Fig. 1a)**. We screened the culture conditions using HCI010 and then validated the results using the other models. HCI010 tumors were harvested from PDX mice after 2-5 months of growth *in vivo*, digested to single cell suspensions and plated into four culture conditions: i) ULA plates and EpiCult (ULA-E); ii) ULA plates and MEGM (ULA-M); iii) Matrigel and EpiCult (MAT-E); and iv) Matrigel with MEGM (MAT-M) **(Fig. 1a)**. Microscopy 7-14 days later showed ULA-E and MAT-E both produced spheroid structures, while ULA-M and MAT-M produced limited growth **(Fig. 1b)**. Flow cytometry analysis of cell viability by annexin V (aV) and propidium iodide (PI) staining showed that MAT-E produced the highest percentage of aV-PI-viable cells (60.6 ± 2.0%; p<0.0001) **(Fig. 1c, Extended Data Fig 1a)**. ULA-E showed 2.3-fold (p<0.0001) lower viability than MAT-E and 1.4-fold (p<0.0001) more aV+PI-cells in early apoptosis **(Fig. 1c, Extended Data Fig 1a)**, indicating MAT-E is superior for producing viable PDX sphere growth.

We next compared the culture conditions for their ability to expand and passage PDX cells. Live cell number was determined before plating and after nine days of culture. Cell counts showed that MAT-E produced a 2.2-fold cell expansion, which was greater than ULA-E (p=0.014) and the other conditions that produced minimal or negative expansion **(Fig. 1d)**. Passaging experiments showed MAT-E could also support extended viable growth of PDX cells. First generation (P0) spheres were harvested, dissociated and re-plated for an additional 14-17 days to produce P1 spheres **(Fig. 1a)**. Flow cytometry analysis of P1 cells showed that MAT-E produced 4-fold greater viability than MAT-M (p<0.0001) **(Fig. 1e)**. Cell counts showed that MAT-E produced a 2-fold expansion of viable cells, which was greater than MAT-M (p=0.008) that reduced total cell number **(Fig. 1f)**.

Further evaluation of MAT-E showed it is sufficient to support growth of other PDX models. HCI001, HCI002 and HCI011 tumors were harvested from PDX mice, dissociated to single cell suspensions and plated in MAT-E conditions. Microscopy analysis showed clear outgrowth of spheres from all three models **(Extended Data Fig 1b)**. We also confirmed that spheres were generated from PDX tumor cells and not contaminating mouse epithelium using a flow cytometry assay previously developed by our laboratory^23^. Spheres generated from each PDX model were dissociated and stained with species-specific antibodies for CD298 (human) and MHC-I (mouse). Flow cytometry analysis showed that >90% of cells were CD298^+^MHC-I^−^human tumor cells **(Extended Data Fig 1c)**. These data show that MAT-E is superior for the viable growth and expansion of human PDX tumor cells *in vitro*. **Extended Data Fig 1d** and **Supplementary videos 1-3** show the kinetics of sphere growth in MAT-E conditions using time lapse imaging.

### PDX tumor sphere cells maintain their global transcriptome program

To determine whether PDX cells maintain their native state following MAT-E culture, we compared the global transcriptome profiles of cultured and uncultured cells by RNA sequencing **(Fig. 2a)**. Cells from HCI002 (n=3) and HCI010 (n=3) tumors were dissociated and split into two matched groups, one for culture in MAT-E (cultured 1-3) and the other for immediate library preparation (uncultured 1-3) **(Fig. 2a)**. CD298^+^MHC-I^−^ cells from both groups were sorted by flow cytometry to enrich for human tumor cells and sequenced at 35 million paired end reads per sample. Reads were aligned to the human hg38 reference genome, and differentially expressed genes were identified using *DESeq2*^24^.

**Figure 2.**
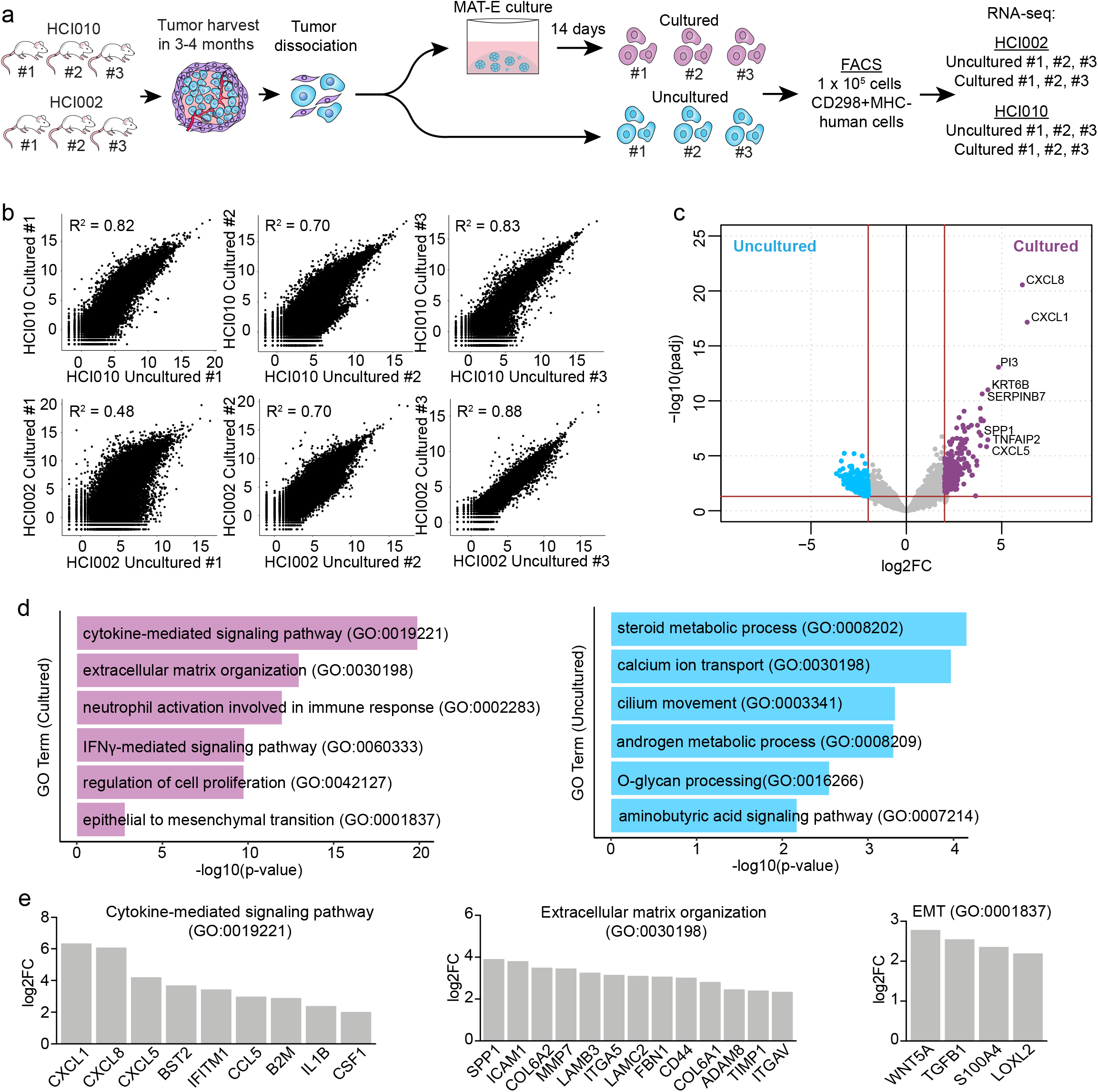
PDX tumor sphere cells maintain their global transcriptome programme. **(a)** Workflow schematic for generation of RNA sequencing dataset to compare transcriptome signatures in paired cultured and uncultured cells from HCI002 and HCI010. **(b)** Scatter plots show correlation of gene expression across the entire transcriptome in paired cultured and uncultured samples (n=22,446 genes). Pearson’s correlation coefficient (R^2^) is shown for each pair. **(c)** Volcano plot shows genes significantly upregulated in cultured (n=277 genes) or uncultured (n=1455 genes) cells that are conserved in all pairs (log2FC>2, adj p<0.05). See **Supplementary Table 1** for full gene list. FC=Fold change. Adjusted *P*-value determined in *DESeq2* by Benjamini-Hochberg adjustment of Wald test *P*-values. **(d)** Bar graph shows GO terms associated with genes upregulated in cultured (n=277 genes) and uncultured (n=1455 genes) cells (log2FC>2, adj p<0.05). *P*-value determined using Enrichr by Fisher’s exact test. **(e)** Bar graphs show select GO term-associated genes upregulated in cultured cells. See **Supplementary Table 1** for full gene list. FC=Fold change. *P*-value determined by Wald test.

Gene expression was compared in paired cultured and uncultured samples from each PDX model by Pearson’s correlation test **(Fig. 2b).** This showed that gene expression was strongly correlated in most pairs, indicating that culture does not substantially alter cell state **(Fig. 2b)**. Differential expression analysis identified 1,732 genes up and downregulated following MAT-E culture that were conserved across all pairs (logFC>2.0, p<0.05) **(Fig. 2c, Supplementary Table 1)**. This represents only 7.7% of the total transcriptome, further indicating that culture does not substantially alter cell state. Notably, canonical breast cell differentiation genes and molecular markers (KRT8, KRT18, KRT14, TP63, ERBB2, ESR1, PGR) were not differentially expressed suggesting the cells maintain their differentiation state and tumor subtype **(Supplementary Table 1)**.

Gene Ontology (GO) analysis of the differentially expressed genes revealed several pathways up and downregulated following MAT-E culture **(Fig. 2d).** Pathways significantly upregulated include ‘cytokine-mediated signaling pathway,’ ‘extracellular matrix (ECM) organization,’ ‘interferon-gamma (IFN_γ_)-mediated signaling pathway’ ‘regulation of cell proliferation,’ and ‘epithelial to mesenchymal (EMT) transition’ **(Fig. 2d).** Analysis of the genes within each GO term identified numerous chemokines (CXCL1, CXCL8 CXCL5), cytokines (CSF1, CCL5, IL1β) and IFN-response genes (BST2, B2M, HLA), suggesting that culture induces a stress-related or inflammatory response **(Fig. 2e)**. We also observed upregulation of 23 ECM-related genes, including ICAM1, COL6A2, MMP7, ITGA5 and LAMC2 **(Fig. 2e)**. Interestingly, cultured cells also displayed increased expression of the EMT-associated genes WNT5A, TGFB1, S100A4, and LOXL2, as well as the CSC marker CD44 **(Fig. 2e, Supplementary Table 1)**. Prior work has shown that normal and breast cancer stem cells display EMT features^25,26^. This may therefore indicate that MAT-E culture selects for stem-like cells, which is consistent with prior reports using 3D culture conditions.^27,28^ However, it is also possible that MAT-E culture induces upregulation of these transcriptome programs.

### PDX tumor sphere cells maintain tumorigenic potential and form spontaneous metastasis *in vivo*

Spontaneous metastasis models are often considered more authentic than experimental metastasis models since the cells must progress through each step of the metastatic cascade in order to metastasize^29^. Prior work has shown that HCI002 and HCI010 develop basal-like primary tumors and spontaneous metastasis following orthotopic transplantation^5,23^. We investigated whether PDX cells maintain these capabilities following sphere culture. We cultured and expanded HCI002 and HCI010 cells in MAT-E for two weeks and orthotopically transplanted serial dilutions of cells into NOD/SCID mice **(Fig. 3a)**. Similar to prior reports with uncultured PDX cell transplants^23^, we observed primary tumor growth from nearly 100% of animals transplanted with cultured cells at each dilution **(Fig. 3b, Extended Data Fig. 2a)**.

**Figure 3.**
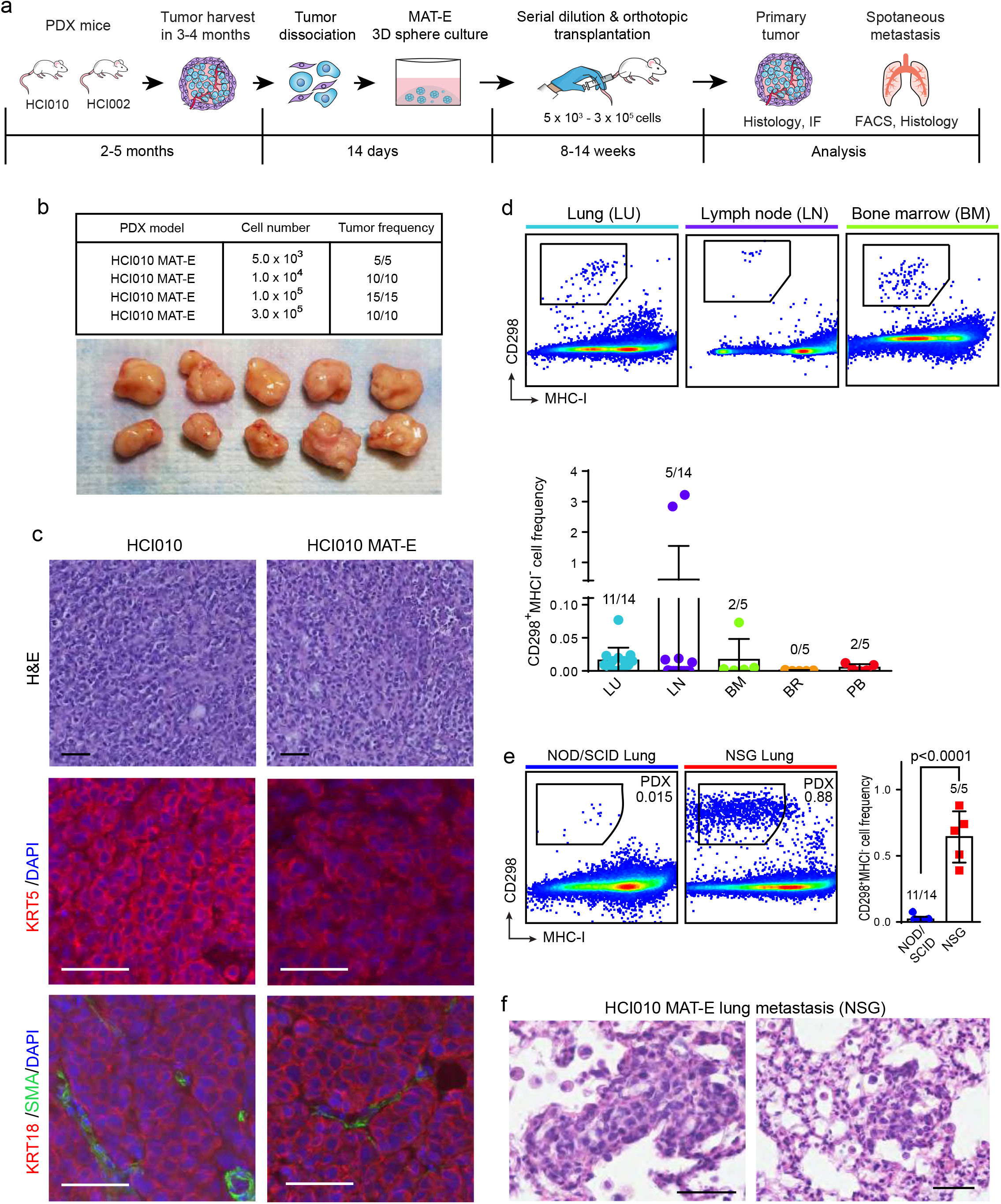
PDX tumor sphere cells maintain tumorigenic potential and form spontaneous metastasis *in vivo*. **(a)** Schematic overview of workflow and timeline for assessing the capacity of cultured PDX cells to generate primary tumors and spontaneous metastasis following orthotopic transplantation. **(b)** Serial dilution and transplantation analysis to determine tumorigenic capacity of cultured cells. HCI010 cells were cultured in MAT-E conditions and injected orthotopically into NOD/SCID mice at increasing dilution (5.0×10^3^ – 3.0×10^5^). Table (top) shows the frequency of tumors generated at each dilution. Representative images (bottom) show primary tumors generated from orthotopic transplantation of 1 × 10^5^ cells. Scale bar = 1cm. **(c)** Histopathological and molecular marker analysis of tumors generated from uncultured (HCI010) and cultured (HCI010 MAT-E) cells. Top panels show representative histopathological appearance of tumor sections stained with hematoxylin and eosin (H&E). Scale bar = 50 µm. Middle and bottom panels show representative images of IF staining for basal (KRT5), luminal (KRT18) and myoepithelial (SMA) cell markers. Scale bar = 50 µm. **(d)** Quantification of spontaneous metastasis in animals transplanted with 1 × 10^5^ cultured HCI010 cells. Representative plots (top) show quantification of CD298^+^MHC-I^−^ human metastatic PDX cells in distal tissues by flow cytometry. Bar graph (bottom) shows quantification of frequency of metastatic cells in a cohort of transplanted animals (n=5-14). Fractions indicate the number of tissues with metastasis, defined by >0.005% CD298^+^MHC-I^−^ cells. Data is represented as the mean ± s.d. LU = Lung, LN = Lymph node, BM = bone marrow, BR = brain, PB = peripheral blood. **(e)** Comparison of lung metastatic burden in NOD/SCID and NSG animals transplanted orthotopically with 1×10^5^ cultured HCI010 cells. Representative plots show CD298^+^MHC-I^−^ human metastatic PDX cells in the lungs 8 weeks after transplant by flow cytometry. Bar graph shows quantification of frequency of metastatic cells in a cohort of transplanted animals. *P*-value determined by unpaired t-test. Data is represented as the mean ± s.d. Fractions indicate the number of tissues with metastasis, defined by >0.005% CD298^+^MHC-I^−^ cells. **(f)** Representative images of metastatic lesions in the lungs of NSG animals transplanted with cultured HCI010 cells identified by H&E staining and histopathological analysis. Scale bar= 50 µm.

We next compared histopathological and molecular features of MAT-E-cultured and native PDX tumors **(Fig. 3c, Extended Data Fig 2b)**. All HCI010 tumors were poorly vascularized, necrotic and showed no epithelioid differentiation, with little nodularity or ductal structure **(Fig. 3c)**. HCI002 tumors showed scant intervening myxoid stroma and thin walled vessels, less necrosis, and retained some small areas of nodular aggregates but were mostly undifferentiated **(Extended Data Fig 2b)**. We observed no pathological differences between native and cultured tumors. Immunofluorescence (IF) staining for canonical basal (KRT5), myoepithelial (SMA) and luminal (KRT18) cell markers further indicated that HCI010 MAT-E tumors maintain their native differentiation state **(Fig. 3c)**. Tumors from both conditions displayed widespread KRT5 and KRT18 expression, and limited SMA expression, as expected for basal-like tumors **(Fig. 3c)**.

We further evaluated the capacity of orthotopic tumors to produce spontaneous metastasis. Flow cytometry analysis identified CD298^+^MHC-I^−^ human metastatic cells in the lungs (11/14), lymph nodes (5/14), bone marrow (2/5), and peripheral blood (2/5) but not the brains (0/5) of transplanted animals **(Fig. 3d).** Metastatic burden was low in most tissues, which is consistent with prior reports **(Fig. 3d,e)**^23^. We therefore tested the capacity of cultured cells to produce spontaneous metastasis in NOD scid gamma (NSG) mice, which lack natural killer (NK) cells and support greater human cell engraftment.^23^ Remarkably, NSG mice displayed >32-fold higher metastatic burden in the lungs than NOD/SCID mice (p<0.0001) **(Fig. 3e)**, and metastatic lesions were detectable by histopathological analysis **(Fig. 3f)**. This is consistent with reports demonstrating the importance of NK cells in controlling metastasis, and shows that different immunodeficient strains can be used to achieve specific experimental contexts^30^. Importantly, we also observed spontaneous metastasis from HCI002 transplants, which displayed lung metastasis (5/5 mice) and circulating tumor cells in the peripheral blood (3/5 mice) **(Extended Data Fig. 2c,d)**. These data show that PDX cells maintain their tumorigenic and metastatic potential after sphere culture, and highlight the utility of our culture-transplant system for studies of spontaneous metastasis from human patient cells.

### PDX tumor sphere cells can be genetically engineered for functional studies of metastasis *in vivo*

The ability to genetically perturb genes of interest is critical for functional investigations of metastasis. We investigated whether PDX tumor sphere cells can be genetically engineered and transplanted to generate spontaneous metastasis *in vivo*. Using a GFP lentiviral construct, we tested the infection efficiency of HCI002 and HCI010 sphere cells grown in MAT-E conditions **(Fig. 4a)**. PDX cells were grown for 2-3 weeks to generate P0 spheres and then harvested and dissociated to generate cell suspensions. P0 cell suspensions were transduced using a combined centrifugation and suspension infection protocol (see Methods), and re-plated in MAT-E conditions for 2-3 weeks to expand and generate P1 spheres **(Fig. 4a)**. Fluorescence microscopy revealed robust GFP expression both within and between P1 spheres **(Fig. 4b,c)**. Quantification of infection efficiency by flow cytometry showed successful transduction with 75.2 ± 6.4% and 23.2 ± 7.6% of HCI010 and HCI002 cells positive for GFP, respectively **(Fig. 4d)**. We next determined whether transduced P1 sphere cells could retain GFP signal and produce spontaneous metastasis following orthotopic transplantation *in vivo*. Flow cytometry sorted GFP^+^ HCI002 and HCI010 P1 sphere cells were transplanted, and primary tumors and lungs were collected and analyzed 60 days later. Whole mount imaging showed that GFP expression was retained in primary tumors **(Fig 4e)**, and quantification by flow cytometry showed that >90% of CD298^+^MHC-I^−^ human PDX tumor cells retained GFP signal **(Fig 4f)**. Analysis of lung tissue sections showed the presence of GFP positive metastatic lesions **(Fig 4g),** and flow cytometry analysis of lung cell suspensions identified CD298^+^MHC-I^−^ GFP^+^ human metastatic cells **(Fig 4h)**.

**Figure 4.**
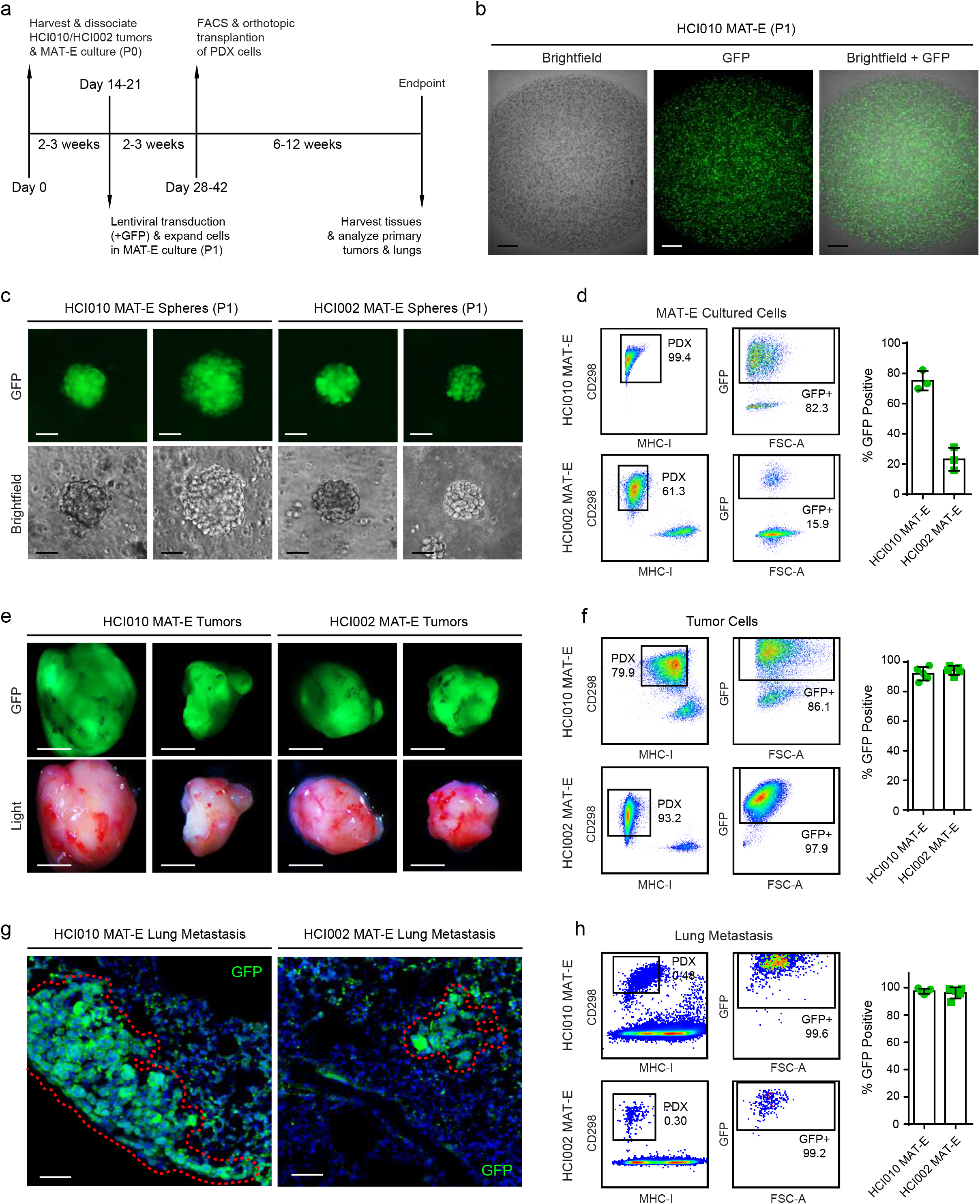
PDX tumor sphere cells can be genetically engineered for functional studies of metastasis *in vivo*. **(a)** Schematic shows workflow for lentiviral engineering of PDX sphere cells *in vitro* followed by analysis of their capacity to form primary tumors and spontaneous metastasis following transplantation *in vivo*. **(b)** Representative images show GFP expression in PDX sphere cultures following lentiviral transduction *in vitro*. HCI010 P0 sphere cells were transduced with GFP lentivirus at MOI=25 and seeded in MAT-E culture at a density of 2 × 10^5^ cells per well. Images show z-stack micrographs of individual wells containing P1 spheres 1 week later. Scale bar= 800 µm. **(c)** Images show GFP expression in representative P1 PDX spheres two weeks post transduction and expansion in MAT-E culture. Scale bar= 50 µm. **(d)** Representative flow cytometry plots show transduction efficiency of P1 PDX sphere cells two weeks post transduction and growth in MAT-E culture. Bar graph quantifies the percentage of PDX cells positive for GFP in each well (n=3). Values represent the mean ± s.d. **(e)** Images show representative PDX tumors produced 60 days post orthotopic transplantation. 1 × 10^5^ CD298^+^MHC-I^−^ GFP^+^ P1 PDX cells were sorted by flow cytometry and transplanted into each mouse (n=5). Scale bar= 5 mm. **(f)** Representative flow cytometry plots show percent of human CD298^+^MHCI^−^ human tumor cells that retain GFP expression following tumor growth *in vivo*. Bar graph quantifies the percentage of human cells positive for GFP in each tumor (n=5). Values are represented as the mean ± s.d. **(g)** Images show representative GFP+ metastatic lesions in the lungs of transplanted animals from (e). Dashed red outline highlights perimeter of lesions. Scale bar= 50 µm. **(h)** Representative flow cytometry plots show percent of human CD298+MHCI-human metastatic cells in the lung that retain GFP expression. Bar graph quantifies the percentage of human cells positive for GFP in each lung (n=5). Values are represented as the mean ± s.d.

Remarkably, >95% of CD298^+^MHC-I^−^ human metastatic cells in the lung also retained GFP expression **(Fig 4h)**. Thus, we demonstrate PDX sphere cells can be robustly engineered, maintain their capacity to form primary tumors and spontaneous metastasis, and retain their genetic alterations *in vivo*.

### PDX tumor sphere cells produce robust experimental metastasis *in vivo*

In experimental metastasis models, cells are injected directly into the circulation via i.v. or i.c. delivery. I.v. injection introduces cells into the venous circulation and favors lung metastasis, while i.c. injection delivers cells into arterial circulation and supports bone and brain metastasis^17,31,32^. These approaches produce metastasis with more robust, rapid and reproducible kinetics than spontaneous models and enable direct investigation of later steps in the metastatic cascade, such as survival in the blood stream, extravasation and seeding in the distal tissue. We investigated whether PDX cells expanded in culture will metastasize following i.v. or i.c. delivery *in vivo*.

We harvested HCI010 and HCI002 tumors from PDX mice and cultured the cells for two weeks in MAT-E conditions **(Fig. 5a)**. 5×10^5^ P0 sphere cells were injected i.v. or i.c. into NOD/SCID mice and peripheral tissues were harvested and analyzed by flow cytometry and histology eight weeks later **(Fig. 5a)**. In i.c. injected animals, we observed CD298^+^MHC-I^−^ human HCI010 cells in the lungs (4/5 mice), bone marrow (3/5 mice), brain (5/5 mice) and peripheral blood (5/5 mice), but not the lymph nodes (0/5 mice) **(Fig. 5b)**. Metastatic burden was remarkably robust in the brain, where HCI010 cells constituted >50% of live cells in three of the mice **(Fig. 5b)**. We also observed metastasis in the lungs, brain and peripheral blood of animals injected i.c. with HCI002 cells **(Extended Data Fig. 3a)**. Like HCI010, metastasis was most robust in the brain, demonstrating the utility of our culture-transplant system for studies of brain metastasis using patient tumor cells.

**Figure 5.**
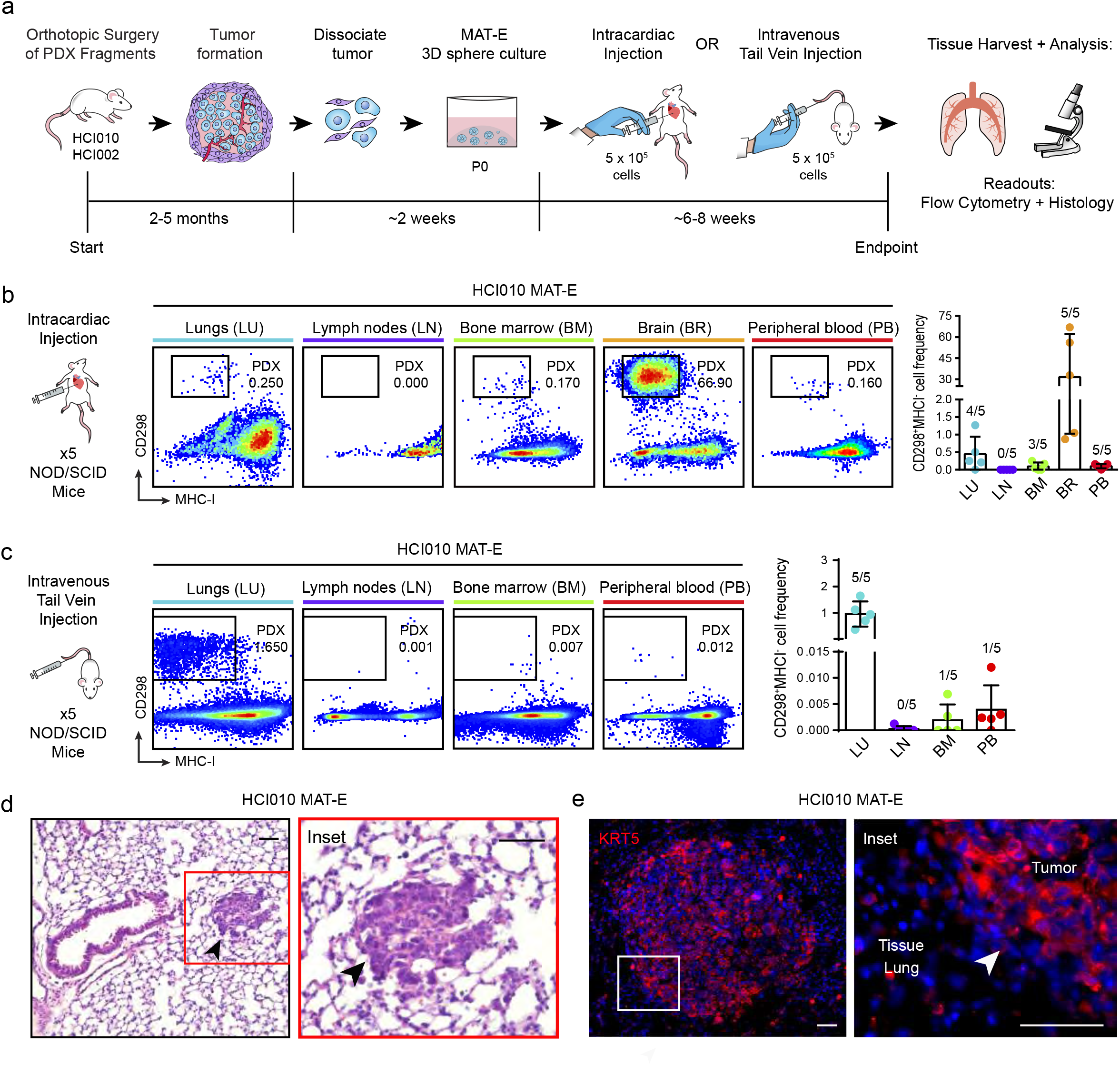
PDX tumor sphere cells produce robust experimental metastasis *in vivo*. **(a)** Schematic overview of workflow and timeline for assessing the capacity of cultured PDX cells to metastasize using i.c and i.v. experimental metastasis models. **(b)** Analysis of metastatic spread following i.c. injection of cultured PDX cells *in vivo*. 5 × 10^5^ cultured HCI010 cells were injected i.c. into NOD/SCID animals and organs were harvested eight weeks later. Flow cytometry plots show human CD298^+^MHC-I^−^ human metastatic cells in the lungs, lymph nodes, bone marrow, peripheral brain, and blood of representative animals (left panels). Bar graph shows quantification of frequency of metastatic cells in a cohort of transplanted animals (n=5) (right panel). Fractions indicate the number of tissues with metastasis, defined by >0.005% CD298^+^MHC-I^−^ cells. Data is represented as the mean ± s.d. **(c)** Analysis of metastatic spread following i.v. injection of cultured PDX cells *in vivo.* 5 × 10^5^ cultured HCI010 cells were injected i.v. into NOD/SCID animals and organs were harvested eight weeks later. Flow cytometry plots show human CD298^+^MHC-I^−^ human metastatic cells in the lungs, lymph nodes, bone marrow, and peripheral blood of representative animals (left panels). Bar graph shows quantification of frequency of metastatic cells in a cohort of transplanted animals (n=5) (right panel). Fractions indicate the number of tissues with metastasis, defined by >0.005% CD298^+^MHC-I^−^ cells. Data is represented as the mean ± s.d. **(d)** Images show representative metastatic lesions identified by histophathological analysis of H&E stained sections of lung tissue from animals described in (c). Arrows indicate metastatic lesion. Scale bar = 50 µm. **(e)** Images show IF staining for KRT5 in lung metastatic lesions from animals described in (c). Arrows indicate metastatic lesions. Scale bar = 50 µm.

In i.v. injected animals, we observed CD298^+^MHC-I^−^HCI010 human cells in the lungs (5/5 mice), bone marrow (1/5 mice), and peripheral blood (1/5 mice), but not the lymph nodes (0/5 mice) **(Fig. 5c)**. As expected, i.v. delivery yielded the highest frequency and burden in the lungs (0.96 ± 0.48%) **(Fig. 5c)**. Analysis of mice injected i.v. with HCI002 cells showed limited metastasis **(Extended Data Fig. 3b)**. We subsequently tested whether i.v. injection into NSG mice would support greater metastasis of HCI002, since NSG mice supported more robust spontaneous metastasis **(Fig. 3e)**. Indeed, flow cytometry analysis showed 5/5 NSG mice developed lung metastasis, compared to 0/5 NOD/SCID mice **(Extended Data Fig. 3b)**. Histopathological analysis confirmed the presence of metastatic lesions in the lungs, and IF staining showed specific expression of the basal cancer subtype marker KRT5 **(Fig. 5d,e, Extended Data Fig. 3c)**. These data demonstrate that patient tumor cells can be expanded in culture and used in experimental metastasis assays.

### Validation of PDX culture-transplant system for authentic investigations of metastasis *in vivo*

To investigate whether our culture-transplant system can be used to faithfully study metastasis, we tested whether known pharmacological inhibitors of metastasis have a similar effect on PDX cells post culture. In prior work, we found that mitochondrial oxidative phosphorylation (OXPHOS) is upregulated in spontaneous lung metastases in PDX mice, and that inhibition of OXPHOS with the complex V inhibitor Oligomycin (Oligo) attenuates lung metastasis^17^. However, functional studies were performed using surrogate cell line models, since functional experiments could not be performed with PDX cells. Here, we determined whether OXPHOS is also critical for the metastatic spread of patient tumor cells using our culture-transplant system **(Fig 6a)**.

**Figure 6.**
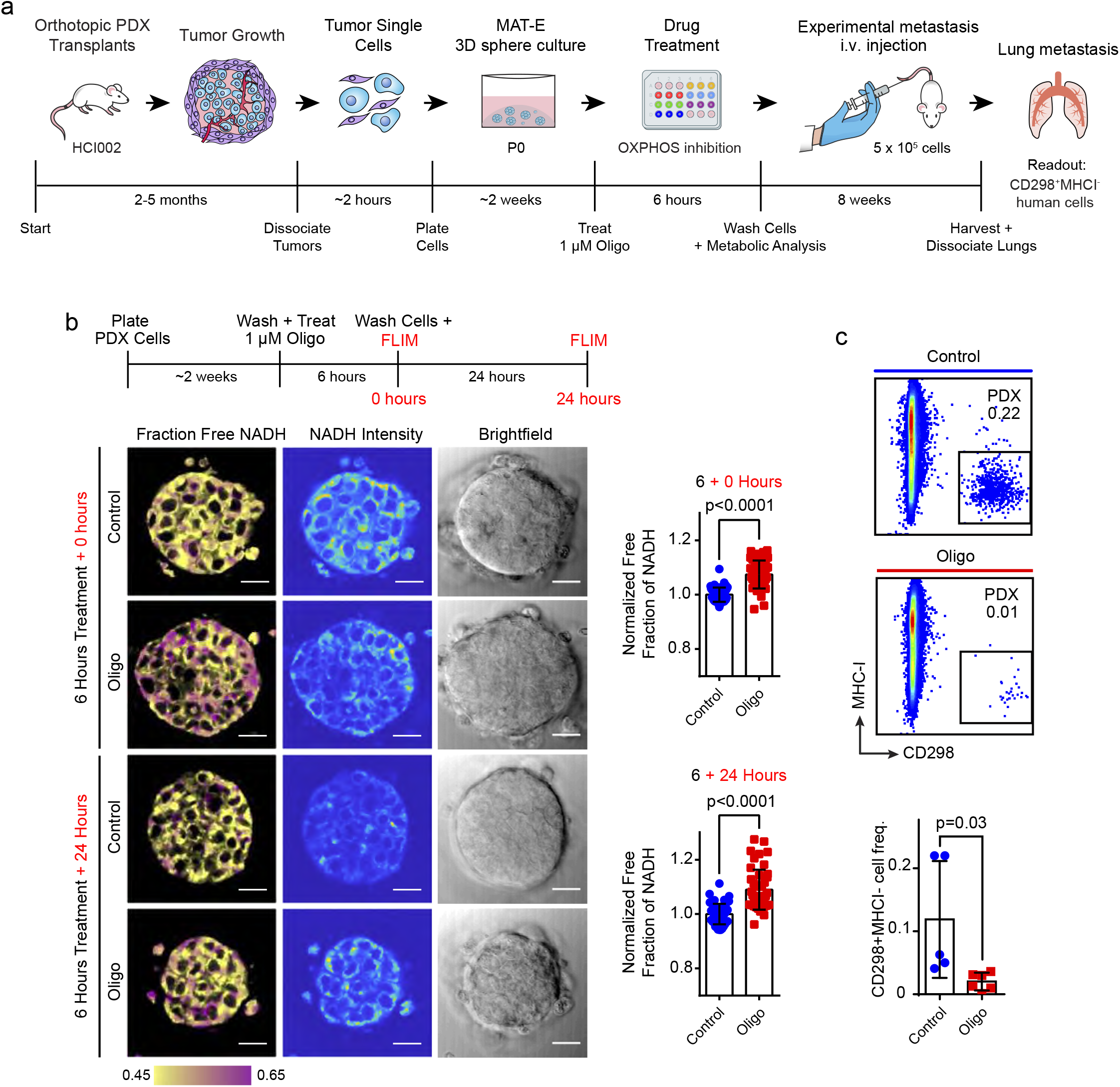
Validation of PDX culture-transplant system for authentic investigations of metastasis *in vivo*. **(a)** Schematic shows experimental workflow to test the effects of OXPHOS inhibition on the metastatic capacity of cultured HCI002 PDX cells following i.v. injection *in vivo*. **(b)** Evaluation of the effects of Oligo treatment on OXPHOS in PDX tumor spheres by FLIM imaging. Schematic shows experimental workflow (top panel). FLIM images show NADH fluorescence signal in representative HCI002 spheres immediately after treatment with 1 µM Oligo or vehicle and replacement with drug-free medium (time = 0 hours) and 24 hours later (+24 hours) (bottom left panels). Scale bar= 20 µm. Bar graphs show quantification of unbound free NADH fraction at 0 and 24 hours (right panels). Values are normalized to the free NADH fraction in control cells at each time point for each replicate, such that a value of 1 represents no change in free NADH compared to control. Values are presented as the mean ± s.d. *P*-value determined by unpaired t-test. Homoscedasticity determined via F-test. Control n= 47 spheres at time t= 0 and 24 hours; Oligo n= 46 spheres at time t= 0 and 24 hours. **(c)** Comparison of metastatic burden in mice transplanted with Oligo vs. vehicle treated PDX cells. 5 × 10^5^ HCI002 PDX sphere cells were treated with 1 µM Oligo or vehicle for 6 hours, washed and transplanted i.v. as described in (a). Flow cytometry plots show the percent of human CD298^+^MHC-I^−^ metastatic cells in the lungs of representative mice eight weeks later. Bar graph shows quantification of frequency of metastatic cells in a cohort of control (n=5) and Oligo (n=6) animals. Values are expressed as mean ± s.d. *P*-value determined by unpaired t-test

We first confirmed that Oligo treatment does not affect PDX cell viability. HCI002 cells were grown in MAT-E culture for two weeks (P0), treated with Oligo for 72 hours and assessed for viability by trypan blue exclusion and aV and PI staining. In contrast to treatment with cytotoxic agents, we observed no overt changes in PDX sphere morphology following Oligo treatment **(Extended Data Fig. 4a)**. Trypan blue exclusion analysis showed there was no significant difference in the percentage of live cells in control (72.3 ± 3.1%) and Oligo (72.3 ± 10.2%) treated conditions (p0.999) **(Extended Data Fig. 4b)**. Analysis of aV and PI by flow cytometry further showed no difference in the percentage of live cells (Oligo, 77.8 ± 1.9%; Control, 81.0 ± 3.1%) (p=0.206) **(Extended Data Fig. 4c,d)**.

We next confirmed that Oligo treatment inhibits OXPHOS in PDX cells using fluorescence lifetime (FLIM) imaging. Previous work has shown that Oligo induces cells to shift from OXPHOS to glycolysis for ATP production^17,33,34^. This shift can be observed using FLIM imaging of NADH, since the fluorescence lifetime of NADH is longer when bound to enzymes involved in OXPHOS (3.4 ns) than when free floating in the cytoplasm during glycolysis (0.4 ns)^35,36^. We treated HCI002 spheres with Oligo for 6 hours, and washed and replaced the media fresh drug-free media **(Fig 6b)**. FLIM showed a 7.4 ± 0.8% increase in the fraction of free NADH at 0 hours (p<0.0001), which was sustained and slightly increased at 24 hours (8.9 ± 0.1%) (p<0.0001) **(Fig. 6b)**, confirming that Oligo induces a shift from OXPHOS to glycolysis in cultured PDX cells.

We determined whether OXPHOS inhibition suppresses the metastatic capacity of cultured PDX cells following i.v. injection. HCI002 P0 sphere cells were treated with Oligo or vehicle control and injected i.v. **(Fig 6a).** Lungs were harvested eight weeks later and the frequency of human CD298^+^MHCI^−^ metastatic cells in each condition was compared by flow cytometry. This revealed a 5.9-fold lower frequency of metastatic cells in mice injected with Oligo (0.02 ± 0.01%, n=5) compared to vehicle (0.12 ± 0.09%, n=6) treated cells (p=0.03) **(Fig. 6c)**. These data validate our culture-transplant system for authentic investigations of metastasis, and show that OXPHOS is critical for metastasis of actual patient tumor cells.

### NME1 promotes lung metastasis from patient tumor cells

We next used our culture-transplant system to investigate potential mechanisms underlying OXPHOS-mediated lung metastasis in PDX animals. In prior work, we found that nucleoside diphosphate kinase A (NME1) is upregulated in spontaneous lung metastases that display increased OXPHOS **(Fig. 7b)**^17^. Increased NME1 expression in primary breast tumors is also associated with worse prognosis in breast cancer patients **(Extended Data Fig 5a)**. NME1 catalyzes the transfer of a phosphate from nucleoside triphosphates (NTPs) to nucleoside diphosphates (NDPs), primarily from ATP to GDP to produce GTP for use in G protein signalling^37^. Several G proteins such as RAC1 an CDC42 promote cell migration and growth and have established pro-metastatic functions^38^. We hypothesized that OXPHOS promotes metastasis in PDX cells through NME1, and tested whether NME1 overexpression promotes metastasis using our culture-transplant system.

**Figure 7.**
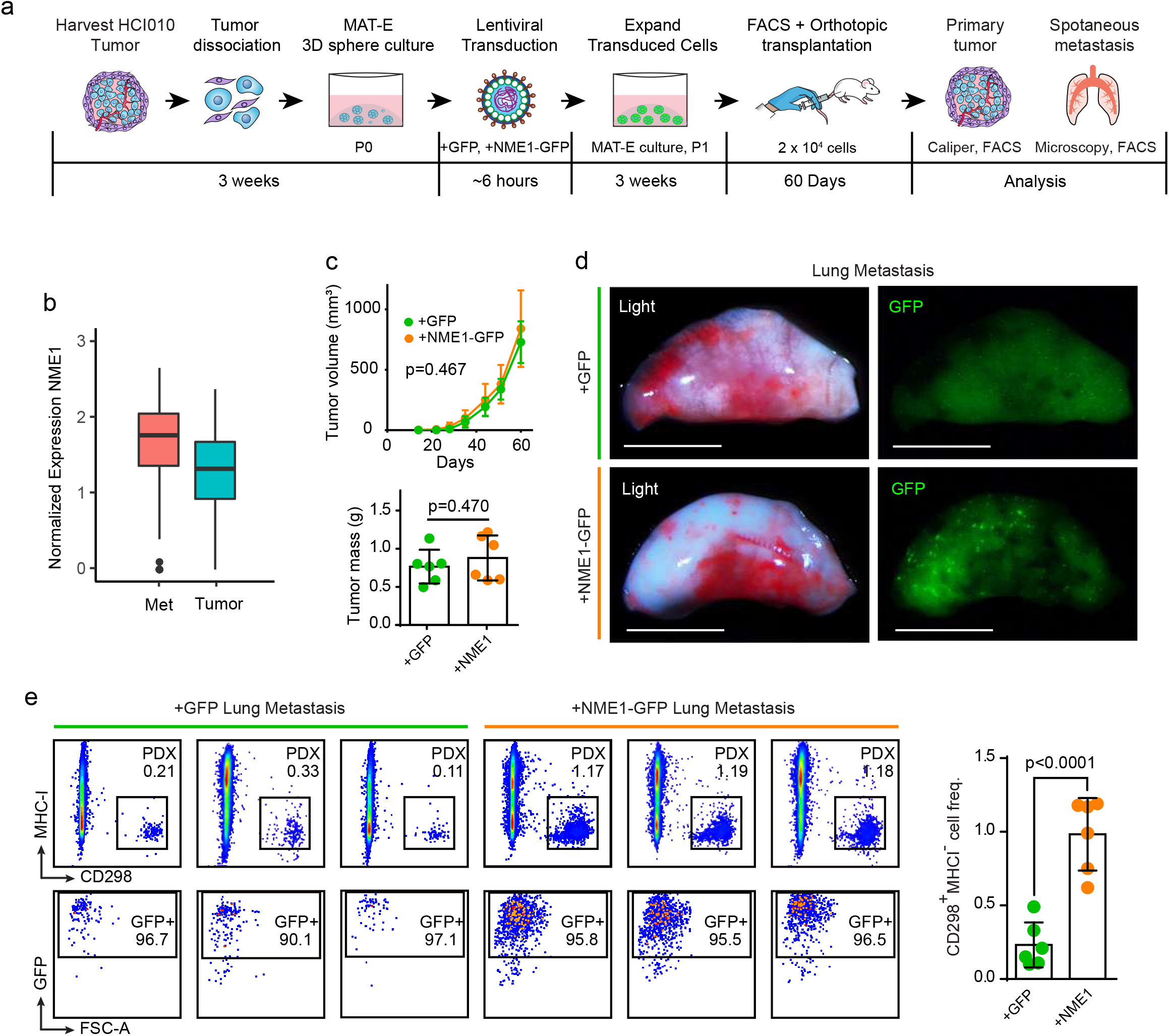
NME1 promotes lung metastasis from patient tumor cells. **(a)** Schematic of experimental workflow to investigate the effect of NME1 overexpression on spontaneous metastasis from cultured PDX tumor cells. **(b)** Box plot shows normalized gene expression of NME1 in metastatic (red, n = 435 cells) and primary tumour cells (blue, n = 684 cells). The lower and upper hinges correspond to the first and third quartiles, and the midline represents the median. The upper and lower whiskers extend from the hinge up to 1.5 * IQR (inter-quartile range). Outlier points are indicated if they extend beyond this range. **(c)** Primary tumor weight and growth kinetics. 2 × 10^4^ HCI010 PDX cells transduced with +NME1-GFP or +GFP lentivirus were transplanted orthotopically and monitored for 60 days (n=6 each condition). Line graph shows tumor volume calculated by caliper measurements (top panel). Bar graph shows tumor weight measured at endpoint (bottom panel). Values are expressed as mean ± s.d. *P*-values determined by unpaired t-test. **(d)** Images show GFP signal identifying metastatic nodules in representative lungs from orthotopically transplanted mice from (c). Scale bar= 5 mm. **(e)** Comparison of lung metastatic burden generated from control versus NME1-overexpressing PDX cells. Flow cytometry plots show the percentage of human metastatic cells in the lungs of three representative mice per group, identified by CD298^+^MHC-I^−^ (top left panels) and GFP (bottom left panels). Bar graph quantifies the frequencies of CD298^+^MHC-I^−^ metastatic cells in the lungs of each animal in the cohort. Values are expressed as mean ± s.d. *P*-value determined by unpaired t-test.

HCI010 cells were cultured in MAT-E conditions for two weeks (P0) and transduced with lentivirus to overexpress NME1 and GFP (+NME1-GFP) or GFP alone (+GFP) **(Fig. 7a)**. Transduced P0 cells were re-plated and expanded in culture for 3 weeks (P1) **(Extended Data Fig. 5b)**, and human GFP^+^ CD298+MHCI-PDX cells were isolated by flow cytometry and transplanted orthotopically into recipient mice for 60 days **(Extended Data Fig. 5c).** qPCR analysis for NME1 showed 3.9-fold increased expression in +NME1-GFP compared to +GFP transduced cells (p=0.0005), confirming successful overexpression **(Extended Data Fig. 5d)**. Analysis of primary tumor growth showed no difference in the growth kinetics or mass of +NME1-GFP (n=6) and +GFP (n=6) groups **(Fig. 7c)**. However, whole mount fluorescence microscopy analysis of lung tissues showed substantially more GFP signal in the +NME1-GFP group, indicating higher metastatic burden **(Fig. 7d)**. This was confirmed by flow cytometry, which showed a >4-fold increase in the frequency of GFP^+^ CD298^+^MHC-I^−^ human metastatic cells in the lungs of +NME1-GFP (0.98 ± 0.25%) compared to +GFP (0.23 ± 0.15%) control mice **(Fig. 7e)**. These findings reveal a pro-metastatic function for NME1 in breast cancer lung metastasis, and provide proof-of-principle for the value of our PDX culture-transplant system for functional investigations of new genes in metastasis using patient tumor cells.

## DISCUSSION

Despite improvements in outcomes for early stage patients, survival rates for metastatic patients have shown limited improvement and metastasis continues to be the major cause of death for cancer patients. A significant roadblock to progress has been the limited availability of models that faithfully recapitulate the biological complexities of metastasis in patients. Despite recent efforts to build large banks of human PDX models to improve authenticity in cancer research, their application in metastasis has been limited by the technical challenges of culturing patient cells in vitro for subsequent functional studies^39,40^. Here, we develop a robust approach for the viable propagation, engineering and expansion of PDX cells in 3D culture for functional studies of metastasis. Using orthotopic transplantation and RNA sequencing, we show that cultured cells maintain tumorigenic potential and display minimal transcriptome changes, indicating that our method facilitates authentic long-term culture of patient tumor cells. We show PDX cells engineered in vitro display robust capacity for spontaneous and experimental metastasis, demonstrating the value of our culture-transplant system for functional studies of metastasis in diverse contexts. To our knowledge, this represents the first adaptation of PDX models for metastasis research *in vivo,* presenting an important technical advance that may help drive innovation for the development of more efficacious therapeutic strategies tailored specifically for metastatic disease.

In addition to technical advances, our study highlights the critical importance of cellular metabolism in breast cancer metastasis. Despite the historical focus on Warburg metabolism in tumor biology, we and others have recently shown that mitochondrial OXPHOS is critical for metastatic progression. However, these studies were conducted using cell line models^17,41,42^, so it remained unclear to what degree OXPHOS is important for metastasis in breast cancer patients.

Here, we used our culture-transplant system to find that OXPHOS inhibition significantly attenuates the lung metastatic capacity of patient tumor cells. We further investigated the potential role of the metabolic enzyme NME1 in OXPHOS-mediated metastasis^17^. Although prior work in melanoma established NME1 as a metastasis suppressor gene,^43–45^ we recently reported its upregulation in lung metastases that display increased OXPHOS. Given that the main function of NME1 is to transfer phosphate from ATP to produce GTP, we hypothesized that NME1 may use ATP generated through OXPHOS to create GTP for G protein signaling pathways that promote metastasis^37,38^. Using our culture-transplant system, we found that NME1 overexpression results in a 4-fold increase in spontaneous lung metastasis. These results reveal a new pro-metastatic role for NME1 in breast cancer, suggesting its function is tissue-context specific, and open the door to future studies investigating the mechanistic link between NME1 and OXPHOS and their potential as therapeutic targets for metastatic disease. Our study also provides key proof-of-principle for the value of our PDX culture-transplant system for functional investigations of new genes in metastasis using patient tumor cells.

## Materials and Methods

### Harvesting and processing tumors into single cells

After 3-6 months of growth, tumors were collected from mice and mechanically dissociated, followed by 2 mg/mL collagenase (Sigma-Aldrich, cat. no. C5138-1G) digestion in medium (DMEM F-12 medium with 5% FBS) at 37°C for 45 minutes on a standard shaker. The digested tumor was washed with PBS, incubated with trypsin (Corning, cat. no. 25-052-Cl) for 10 minutes at 37°C, washed again, and then treated with DNaseI (Worthington Biochemical, cat. no. LS002139). The cells were filtered through a 100 µm strainer. Live cell concentration was checked using the Countess II automated cell counter (ThermoFisher Scientific Inc., Carlsbad, CA, USA).

### Culturing PDX cells

PDX tumors were dissociated into single cells and plated in various culture conditions. PDX cells were plated in 6-well ULA plates (Fisher Scientific, cat. no. 07-200-601) at 2.5 × 10^5^ cells/well and topped off with 1 mL of either EpiCult™ Medium (StemCell Technologies, cat. no. 05610) supplemented with 5% FBS, 10 ng/mL human epidermal growth factor, and 10 ng/mL basic fibroblast growth factor or MEGM™ culture medium (Lonza, cat. no. CC-3150), completed as per manufacturer’s protocol. Medium was changed when necessary.

PDX single cells were also cultured in standard flat bottom 24-well plates. The cells were plated at a seeding density of 1.0 × 10^5^ to 2.5 × 10^5^ cells/well in a 1:1 mix of Matrigel (growth factor reduced) (Corning, cat. no. 356231) and EpiCult™ or MEGM™. 1 mL of EpiCult medium was added to each well, and changed as necessary. Cells were harvested anytime between day 9-21 using dispase (StemCell Technologies, cat. no. 07913) and trypsin to dissociate Matrigel and spheres. All PDX cells from all culture conditions were grown at 37°C and 5% CO_2_.

### Generation of RNA sequencing dataset

We generated a dataset of 3 biological replicates of paired HCI010 uncultured and cultured cells, and paired HCI002 uncultured and cultured cells. We harvested and dissociated HCI010 and HCI002 PDX tumors using methods described above. We split our single cell suspension in half and stained half of the uncultured HCI010 and HCI002 cells for FACS using fluorescent labeled antibodies for human specific CD298 (Biolegend, cat. no. 341704) and mouse-specific MHC-I (eBioscience, cat. no. 17-5957-82). 1.0×10^5^ uncultured cells were sorted for each patient for RNA isolation. The other half of the cell suspension was used for culturing. HCI010 and HCI002 PDX cells were plated in 24-well plates at a seeding density of 2.5 × 10^5^ cells/well and topped off with 1 mL of EpiCult. Cells were grown in culture for 2 weeks, and then harvested using dispase and trypsin. Cultured cells were stained for FACS using fluorescent labeled antibodies for human-specific CD298 and mouse-specific MHC-I. 1.0×10^5^ cultured cells were sorted for each patient. After sorting, uncultured and cultured cells were processed for RNA extraction, followed by cDNA synthesis and amplification. Library construction was performed with the Takara Clontech SMARTer Stranded Total RNA-seq kit v2. The libraries were sequenced at a depth of 35 million paired end reads for each sample on the NovaSeq 6000. RNA sequencing data is available on the GEO database under the accession code GSE153887.

### Processing and analysis of bulk RNA-seq data

Quality of reads was accessed using FastQC software. Reads were aligned to reference genome GRCh38/hg38 from UCSC using Spliced Transcripts Alignment to a Reference (STAR) software. The un-normalized count matrix was filtered to exclude any genes with 0 reads, and we designated the uncultured samples as the reference level. We then performed differential gene expression analysis using DESeq2 1.28.1 in R 4.0.0.

### Mouse strains

NOD/SCID and NSG mice were purchased from The Jackson Laboratory (Bar Harbor, Maine, USA). All mice were maintained in a pathogen-free facility. All mouse procedures were approved by the University of California, Irvine, Institutional Animal Care and Use Committee.

### Orthotopic transplantation and i.c and i.v injection models of cultured PDX cells for modeling metastasis

PDX cells were harvested after two to three weeks of culture in Matrigel and EpiCult. Dispase and trypsin were used to dissociate Matrigel and spheres to single cells. For orthotopic transplants, 2.0 × 10^4^ − 1.0 × 10^5^ cells were suspended in 15-20 µL of a 1:1 mixture of PBS and Matrigel and injected into the number 4 mammary fat pads of NOD/SCID and NSG mice using a 25 µL Hamilton syringe. For experimental metastasis models, cells were suspended at a concentration of 5×10^6^ cells/mL. 5×10^5^ cells in 100ul of PBS were injected into the right ventricle of each NOD/SCID or NSG mouse for i.c. injections or into the tail vein for i.v. injections.

### Analysis of primary tumors and metastasis in distal tissues

Following 8-14 weeks post orthotopic transplants, or ~8 weeks post i.c. or i.v. injections, primary tumors and peripheral tissues including lungs, lymph nodes, bone marrow, peripheral blood, and brains were harvested, dissociated to single cells and stained with fluorescent conjugated antibodies for CD298 (Biolegend, cat. no. 341704) and MHC-I (eBioscience, cat. no. 17-5957-82), and flow cytometry was used to analyze and quantify disseminated CD298^+^MHC-I^−^ PDX cells using the BD FACSAria Fusion cell sorter (Becton, Dickinson and Company, Franklin Lakes, NJ, USA) as described previously^23^. Primary tumor volumes were calculated using caliper measurements with the equation: volume of an ellipsoid= 1/2(length × width^2^). Whole mount images of primary tumors and lungs were taken with a Leica MZ10 F modular stereo microscope (Leica Microsystems, Buffalo Grove, IL, USA).

### Histology and pathological analysis

Tissues were harvested from mice. The lungs and tumors were fixed in 4% formaldehyde. After overnight fixation at 4°C, the tissues were processed for paraffin embedding using standard protocols. The paraffin-embedded tissues were cut into 7-μm sections using a Leica microtome, rehydrated, and stained with haematoxylin and eosin. For immunofluorescent staining, slides were subjected to antigen retrieval in a 10 mM citric acid buffer (0.05% Tween 20, pH 6.0). We then blocked non-specific binding with a blocking buffer (0.1% Tween 20 and 10% FBS in PBS) for 20 minutes, and then incubated tissues overnight at 4°C with primary antibodies. Slides were washed with PBS, and then incubated at room temperature with secondary antibodies for an hour, followed by washing with PBS, and mounting with VECTASHIELD Antifade Mounting Medium with DAPI (Vector Laboratories, H-1200-10). Images were taken using a BZ-X700 Keyence microscope (Keyonce Corp. of America, Itasca, IL, USA). Pathological analysis was performed by Dr. Robert A. Edwards, a board-certified pathologist and the director of the Experimental Tissue Resource in the UC Irvine Health Chao Family Comprehensive Cancer Center.

### Lentivirus transduction for genetic engineering of PDX cells

PDX cells were cultured for approximately 2-3 weeks in Matrigel and EpiCult, removed from Matrigel with dispase, dissociated to single cells with trypsin, neutralized with DMEM medium with 10% FBS, and washed with PBS with a 5 min centrifugation at 300 × g. Cells were suspended in premade lentivirus solution at a multiplicity of infection (MOI) of 25 and the final volume was topped off to no greater than 100 µL with Opti-MEM™ I Reduced Serum Medium (ThermoFisher Scientific, Cat No. 31985070). GFP control (+GFP) and NME1 (+NME1) lentiviral expression vectors were packaged into lentiviral particles and purchased from VectorBuilder Inc., Chicago, IL, USA (+GFP Cat. No. VB190812-1255tza, +NME1 Cat. No. VB180802-1083ueg). Cells were subjected to an hour long spinfection at 300 × g at 4°C, resuspended and incubated in the same lentivirus solution for an additional 4-5 hours in 96-well round bottom ultra low attachment plate wells at 37°C and 5% CO_2_. Cells were centrifuged for 5 min at 300 × g, seeded in Matrigel and EpiCult as described above and expanded for an additional 2-3 weeks at 37°C and 5% CO_2_. Medium was replaced as needed. After 2-3 weeks in Matrigel and EpiCult cultured, PDX cells were removed from Matrigel with dispase, dissociated to single cells with trypsin to form single cell suspensions for transplantation into mice. Cell sorting for GFP-positive CD298^+^MHC-I^−^ PDX cells with fluorescently tagged antibodies for CD298 (Biolegend, cat. no. 341704) and MHC-I (eBioscience, cat. no. 17-5957-82) was performed using the BD FACSAria Fusion cell sorter (Becton, Dickinson and Company, Franklin Lakes, NJ, USA).

### Pharmacologic studies

Cells were treated with oligomycin (Oligo) (MP Biomedicals, Cat. No. 0215178610) for the inhibition of OXPHOS and treated with Taxol (Sigma-Aldrich Canada, Cat. No. T7402), staurosporine (STS) (Sigma-Aldrich Canada, Cat. No. S4400), and doxorubicin (DOX) (Sigma-Aldrich Canada, Cat. No. D1515) as positive controls for cell death. All compounds were prepared as concentrated stock solutions in DMSO and cells were treated with diluted working solutions in culture medium at DMSO concentrations no greater than 0.5% (v/v) using drug doses based off previous studies^17,46^.

### Cell death and viability assays

Trypan Blue exclusion assays were performed with the Countess II automated cell counter (ThermoFisher Scientific Inc., Carlsbad, CA, USA). For analysis of cell death, cells were stained with aV-FITC, diluted 1:100 (GeneTex Cat. No. GTX14082) and PI, diluted 1:100 (ThermoFisher Scientific Cat. No. P3566) for 15 minutes in aV binding buffer (10 mM HEPES, 140 mM NaCl, 2.5 mM CaCl_2_, pH 7.4). Fluorescence microscopy was performed with the BZ-X700 Keyence microscope (Keyonce Corp. of America, Itasca, IL, USA), and flow cytometry analysis was performed using the BD FACSAria Fusion cell sorter (Becton, Dickinson and Company, Franklin Lakes, NJ, USA).

### NADH fluorescence lifetime imaging

NADH fluorescence lifetime images were acquired with an LSM 880 confocal microscope (Zeiss) with a 40×1.2 NA C-Apochromat water-immersion objective coupled to an A320 FastFLIM acquisition system (ISS). A Ti:Sapphire laser (Spectra-Physics Mai Tai) with an 80 MHz repetition rate was used for two-photon excitation at 740 nm. The excitation signal was separated from the emission signal by a 690 nm dichroic mirror. The NADH signal was passed through a 460/80 nm bandpass filter and collected with an external photomultiplier tube (H7522P-40, Hamamatsu). Cells were imaged within a stage-top incubator kept at 5% CO_2_ and 37°C. FLIM data was acquired and analyzed with the SimFCS 4 software developed at the Laboratory for Fluorescence Dynamics at UC Irvine. Calibration of the system was performed by acquiring FLIM images of coumarin 6 (~10 µM), which has a known lifetime of 2.4 ns in ethanol, to account for the instrument response function.

### Phasor FLIM NADH fractional analysis

NADH assumes two main physical states, a closed configuration when free in solution, and an open configuration when bound to an enzyme^47^. These two physical states have differing lifetimes, 0.4 ns when in its free configuration, and 3.4 ns when in its bound configuration ^48–50^. To quantify metabolic alterations, we performed fractional analysis of NADH lifetime by calculating individual pixel positions on the phasor plot along the linear trajectory of purely free NADH lifetime (0.4 ns) and purely bound NADH lifetime (3.4 ns). We quantified the fraction of free NADH by calculating the distance of the center of mass of a spheroid’s cytoplasmic NADH FLIM pixel distribution to the position of purely bound NADH divided by the distance between purely free NADH and purely bound NADH on the phasor plot. These segmentation and phasor analysis methods are described in detail elsewhere^51^.

### Single cell analysis

To analyze the differences in the expression of NME1 between metastases and tumors in these models, we accessed our previously published dataset on these PDX models, which included single cell gene expression profiles of metastatic and primary tumor cells^17^. The expression matrices were log-transformed into log[transcripts per kilobase million + 1] matrices and loaded into the Seurat analysis package^52^.

### qPCR analysis

RNA was extracted by using Quick-RNA Microprep Kit (Zymo Research, R1050) following manufacturer’s protocol. RNA concentration and purity were measured with a Pearl nanospectrophotometer (Implen). Quantitative real-time PCR was conducted using PowerUp SYBR green master mix (Thermo Fisher Scientific, A25742) and primer sequences were found in Harvard primer bank and designed from Integrated DNA Technologies; GAPDH forward primer 5’-CTCTCTGCTCCTCCTGTTCGAC −3’, GAPDH reverse primer 5’-TGAGCGATGTGGCTCGGCT −3’; NME1 forward primer 5’-AAGGAGATCGGCTTGTGGTTT −3’, NME1 reverse primer 5’-CTGAGCACAGCTCGTGTAATC −3’. Gene expression was normalized to the GAPDH housekeeping gene. For relative gene expression 2^negΔΔCt values were used. The statistical significance of differences between groups was determined by unpaired t-test using Prism 6 (GraphPad Software, Inc).

### Relapse-free survival analysis

Kaplan-Meier (KM) survival curves were generated to perform relapse-free survival analysis on breast cancer patient primary tumor microarray data (of all breast cancer patients and individual subtypes of breast cancer) from the KM plotter database^53^. All KM plots are displayed using the “Split patients by median” parameter.

## Acknowledgements

We thank Dr. Zena Werb at the University of California, San Francisco for thoughtful discussions and feedback on project overview and study design. We thank Aaron Longworth, Armani Oganyan, Nathan Ryan James, Anh Thien Phung, Scott Nguyen, Lannhi Nguyen and Jennifer Nguyen for their technical assistance and animal handling. We thank Dr. Alana Welm at the Huntsman Cancer Institute for generously providing PDX models. We thank the Genomics High Throughput Facility at the University of California, Irvine for conducting the bulk RNA sequencing of the PDX samples. We wish to acknowledge the support of the Chao Family Comprehensive Cancer Center Experimental Tissue Resource, supported by the National Cancer Institute of the National Institutes of Health under award number P30CA062203.

## Author Contributions

D.A.L and K.K. supervised research. D.A.L, K.K., D.M., and G.A.H. designed research. D.M., G.A.H., A.E.Y.T.L., H.A., K.B., K.R.D., M.R., J.W.W., R.T.D, K.T.E., M.Y.G.M., R.L., R.A.E., M.A.D., K.K., and D.A.L. performed research; G.A.H., K.B., and J.W.W. performed bioinformatic analyses; D.M., G.A.H., K.K. and D.A.L. wrote the paper manuscript, and all authors discussed the results and provided comments and feedback.

## Data availability

The authors declare that all data supporting the findings of this study are available within the article and its supplementary information files or from the corresponding author upon reasonable request.

## Extended Data Figure Legends

**Supplementary Videos 1-3: Growth of HCI010 cells in MAT-E culture conditions.**

Time-lapse imaging of HCI010 PDX cells in MAT-E culture at day 7 post seeding. Growth shown over the course of 48 hours.

**Supplementary Table 1: Identification of genes differentially in PDX cells after culture.** Differential expression analysis identified 1,732 genes up and downregulated following MAT-E culture that were conserved across all PDX models (logFC>2.0, p<0.05). Adjusted *P*-values were determined in *DESeq2* by Benjamini-Hochberg adjustment of Wald test *P*-values.

**Extended Data Figure 1:**
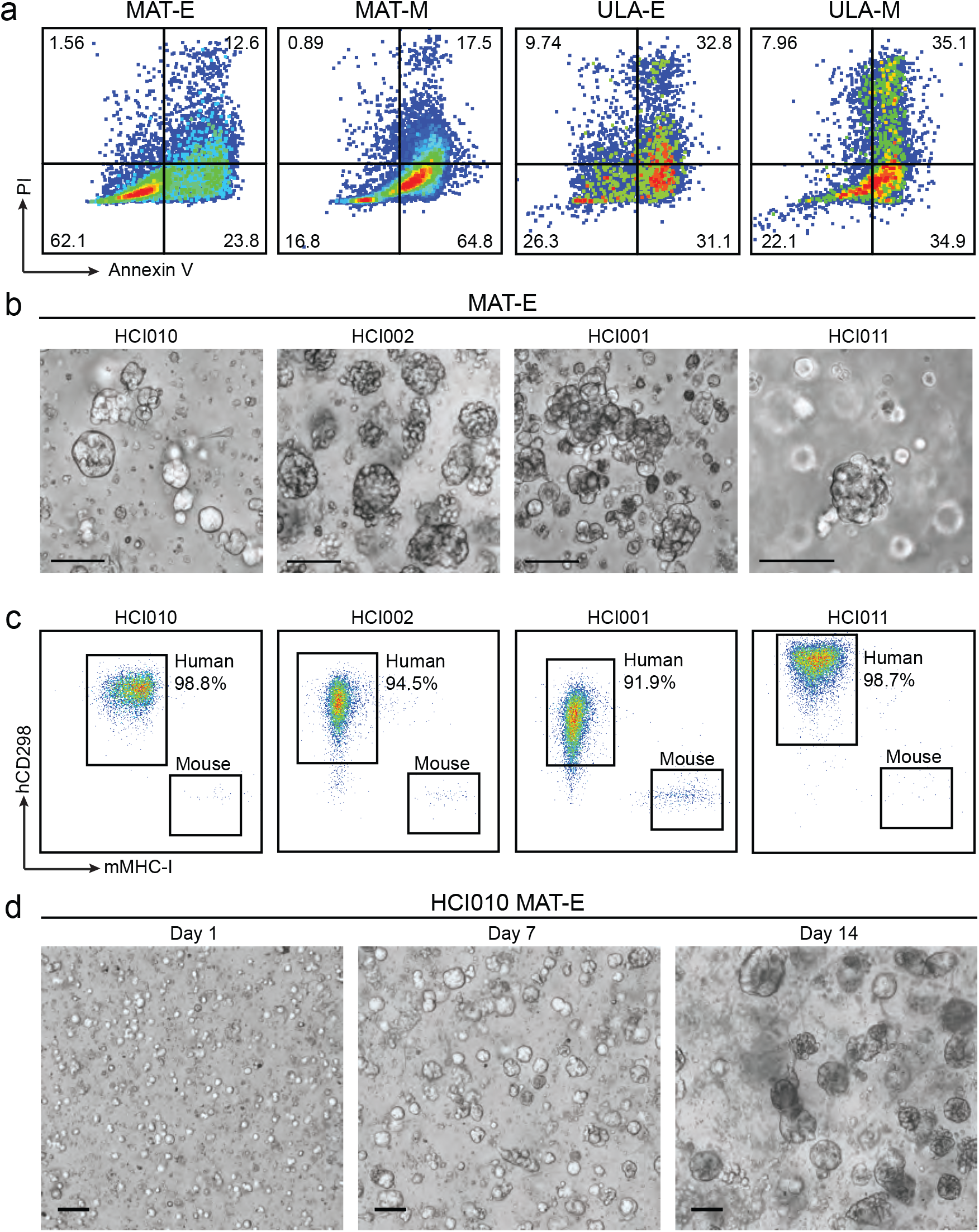
Comparison of PDX cell culture conditions. **(a)** Representative flow cytometry plots show viability of HCI010 cells by aV and PI staining after culture as described in Fig 1a. **(b)** Representative brightfield images show spheres generated by HCI010, HCI002, HCI001 and HCI011 tumor cells grown in MAT-E culture conditions. Scale bar= 100 µm **(c)** Flow cytometry analysis to determine species identity of spheres generated in (b). Representative plots show the percent of human CD298+MHCI-tumor cells in each culture. **(d)** Timelapse imaging of sphere growth from HCI010 cells in MAT-E conditions. Representative brightfield images at day 1, 7, and 14 are shown. See also Supplementary videos 1-3. Scale bar= 100 µm.

**Extended Data Figure 2:**
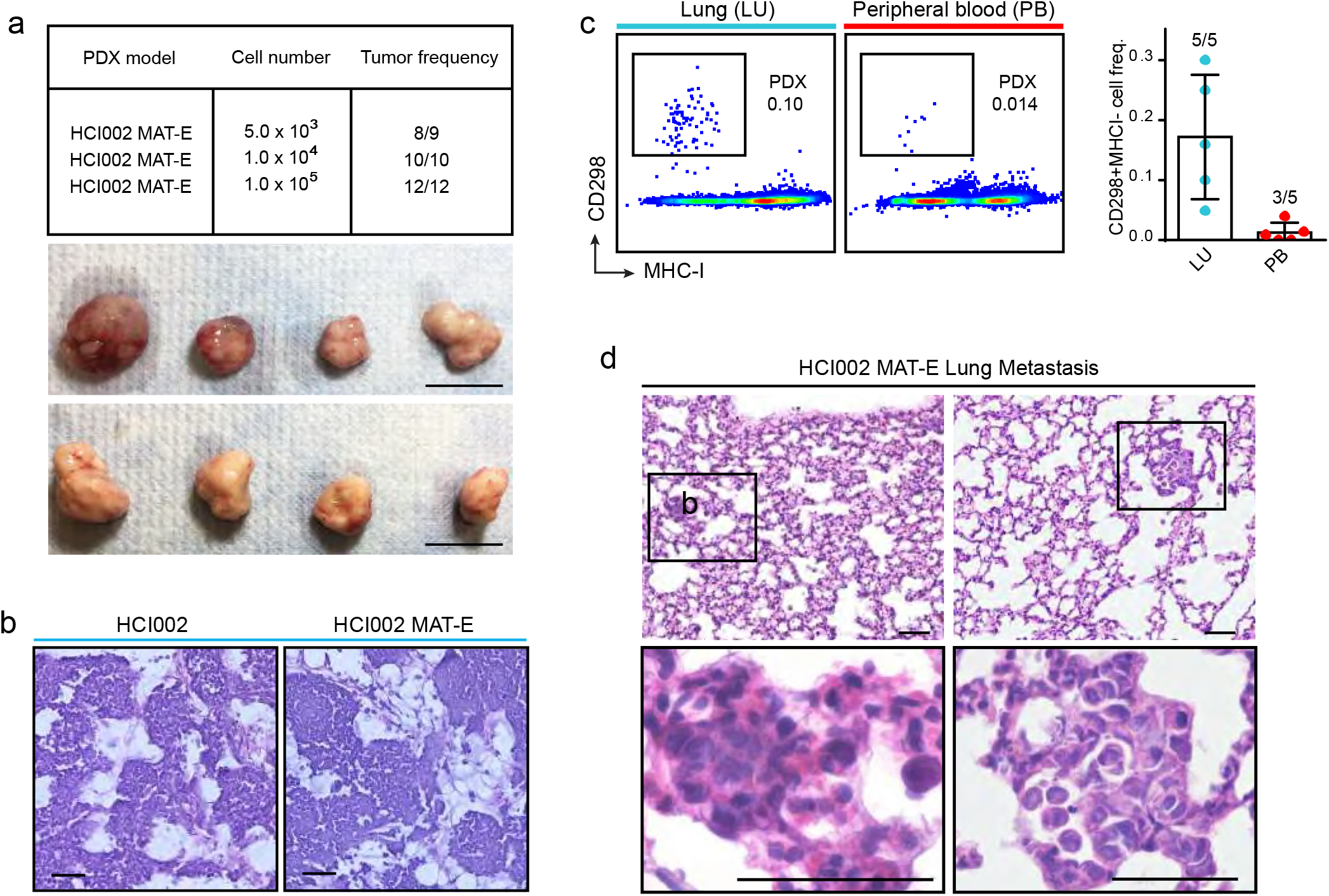
Analysis of primary tumors and spontaneous metastases generated by cultured PDX cells. **(a)** Serial dilution and transplantation analysis to determine tumorigenic capacity of cultured cells. HCI002 cells were cultured in MAT-E conditions and injected orthotopically into NOD/SCID mice at increasing dilution (5.0×10^3^ – 1.0×10^5^). Table (top) shows the frequency of tumors generated at each dilution. Representative images (bottom) show primary tumors generated from orthotopic transplantation of 1 × 10^5^ cells after 12 weeks *in vivo*. Scale bar = 1cm. **(b)** Histopathological analysis of tumors generated from uncultured (HCI002) and cultured (HCI002 MAT-E) cells. Representative images show tumor sections stained with hematoxylin and eosin (H&E). Scale bar = 50 µm. **(c)** Quantification of spontaneous metastasis in animals transplanted with 1 × 10^5^ cultured HCI002 cells. Representative plots (left) show CD298^+^MHC-I^−^ human metastatic PDX cells in the lung and peripheral blood by flow cytometry. Bar graph (right) shows quantification of frequency of metastatic cells in a cohort of transplanted animals (n=5). Fractions indicate the number of tissues with metastasis, defined by >0.005% CD298^+^MHC-I^−^ cells. Data is represented as the mean ± s.d. LU = Lung, PB = peripheral blood. **(d)** Representative images of metastatic lesions in the lungs of animals transplanted with cultured HCI002 cells identified by H&E staining and histopathological analysis. Scale bar= 50 µm.

**Extended Data Figure 3:**
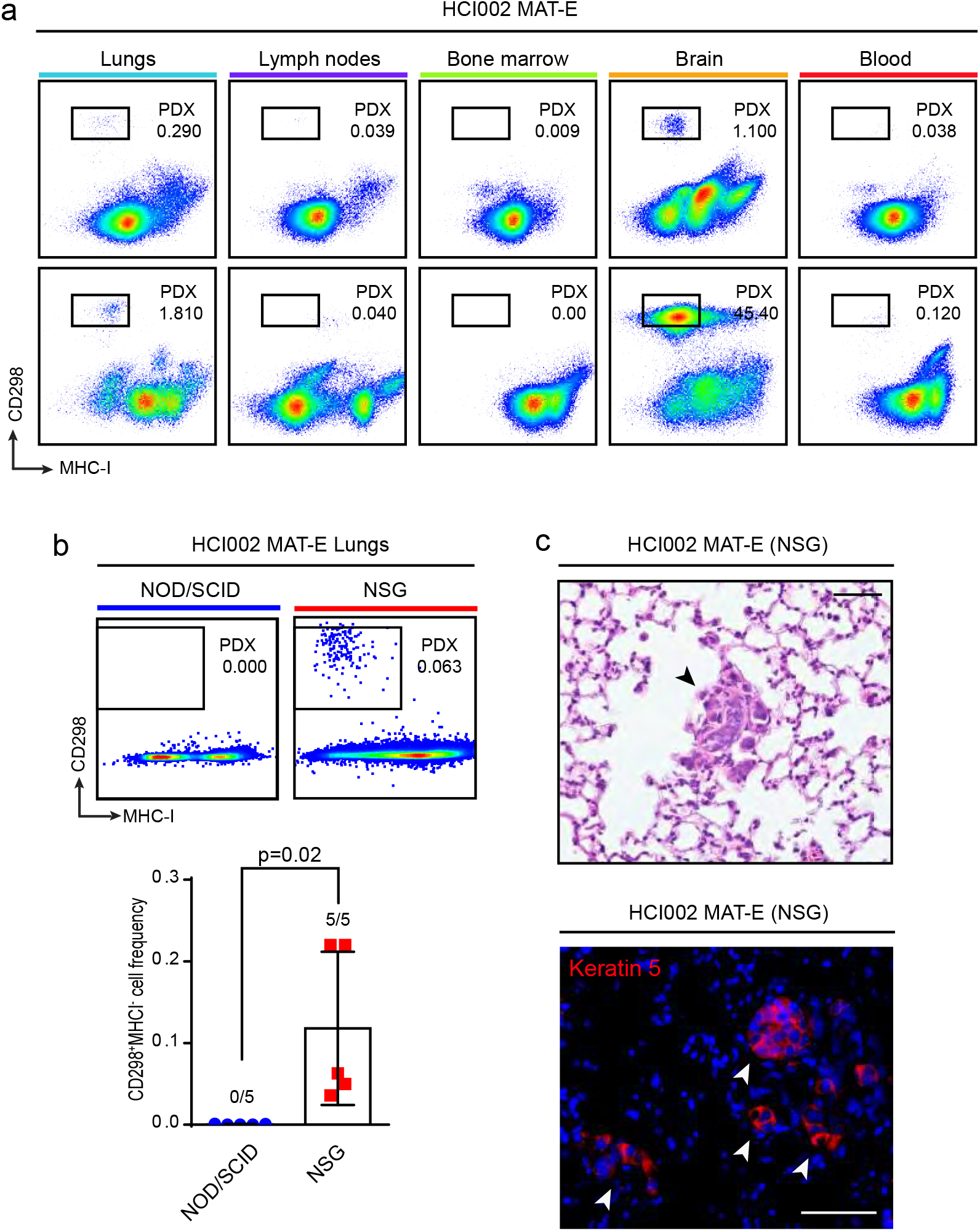
HCI002 tumor spheres produce experimental metastasis *in vivo*. **(a)** Analysis of metastatic spread following i.c. injection of cultured HCI002 cells *in vivo.* Representative flow cytometry plots show CD298^+^MHC-I^−^ human metastatic cells in the lungs, lymph nodes, bone marrow, brain, and blood 8 weeks following i.c. injection of 5 × 10^5^ cultured HCI002 cells into NOD/SCID mice. **(b)** Comparison of experimental metastasis from cultured HCI002 cells in NOD/SCID and NSG mice. Flow cytometry plots show percent of human CD298^+^MHC-I^−^ metastatic cells in the lungs of representative mice 8 weeks following i.v. injection of 5 × 10^5^ HCI002 cultured cells (top plots). Bar graph shows quantification of frequency of metastatic cells in a cohort of transplanted animals (n=5) (bottom). Fractions indicate the number of lungs with metastasis, as defined by >0.005% CD298^+^MHC-I^−^ cells. Data is represented as the mean ± s.d.. *P*-value was determined by unpaired t-test. **(c)** Images show representative metastatic lesions (arrows) in the lungs of NSG mice identified by H&E staining (top) and IF staining for KRT5 (bottom). Scale bar = 50 µm.

**Extended Data Figure 4:**
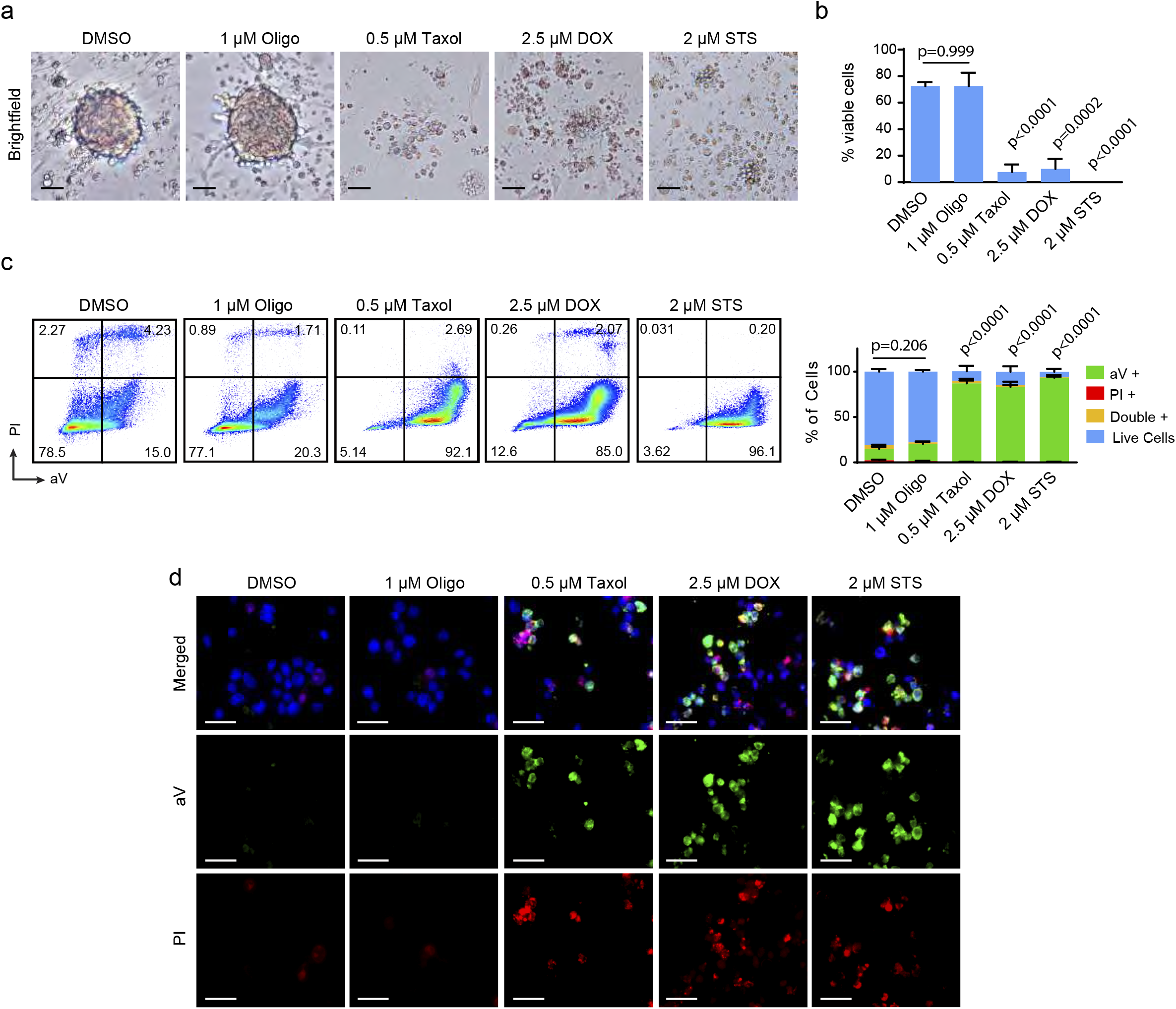
Cytotoxicity and cell death analysis of PDX cells post drug treatment. **(a)** Z-stack brightfield images of representative HCI002 MAT-E spheroids grown for two weeks and treated with the indicated drugs for 72 hours. Scale bar= 50 µm. Oligo= oligomycin; DOX= doxorubicin; STS= staurosporine. **(b)** Trypan blue exclusion viability assay of HCI002 MAT-E cells grown for two weeks and treated with the indicated drugs for 72 hours. Values expressed as mean ± s.d. from triplicate wells (n=3). *P*-values were determined by unpaired t-tests of each condition vs. the DMSO control group. Taxol, DOX and STS were used as positive controls for cell death. **(c)** Representative flow cytometry plots of aV and PI dead cell analysis of HCI002 MAT-E cells treated for 72 hours. Bar graph shows percent of live and dead HCI002 cells by aV and PI. Data represented as mean ± s.d. (n=3). *P*-values were determined by unpaired t-tests of each condition vs. the DMSO control group for the percent of aV-PI-viable cells. Taxol, DOX and STS were used as positive controls for cell death. **(d)** Representative aV and PI fluorescence micrographs of HCI002 MAT-E cells treated for 72 hours. Blue= Hoechst nuclear counterstain. Scale bar= 25 µm.

**Extended Data Figure 5:**
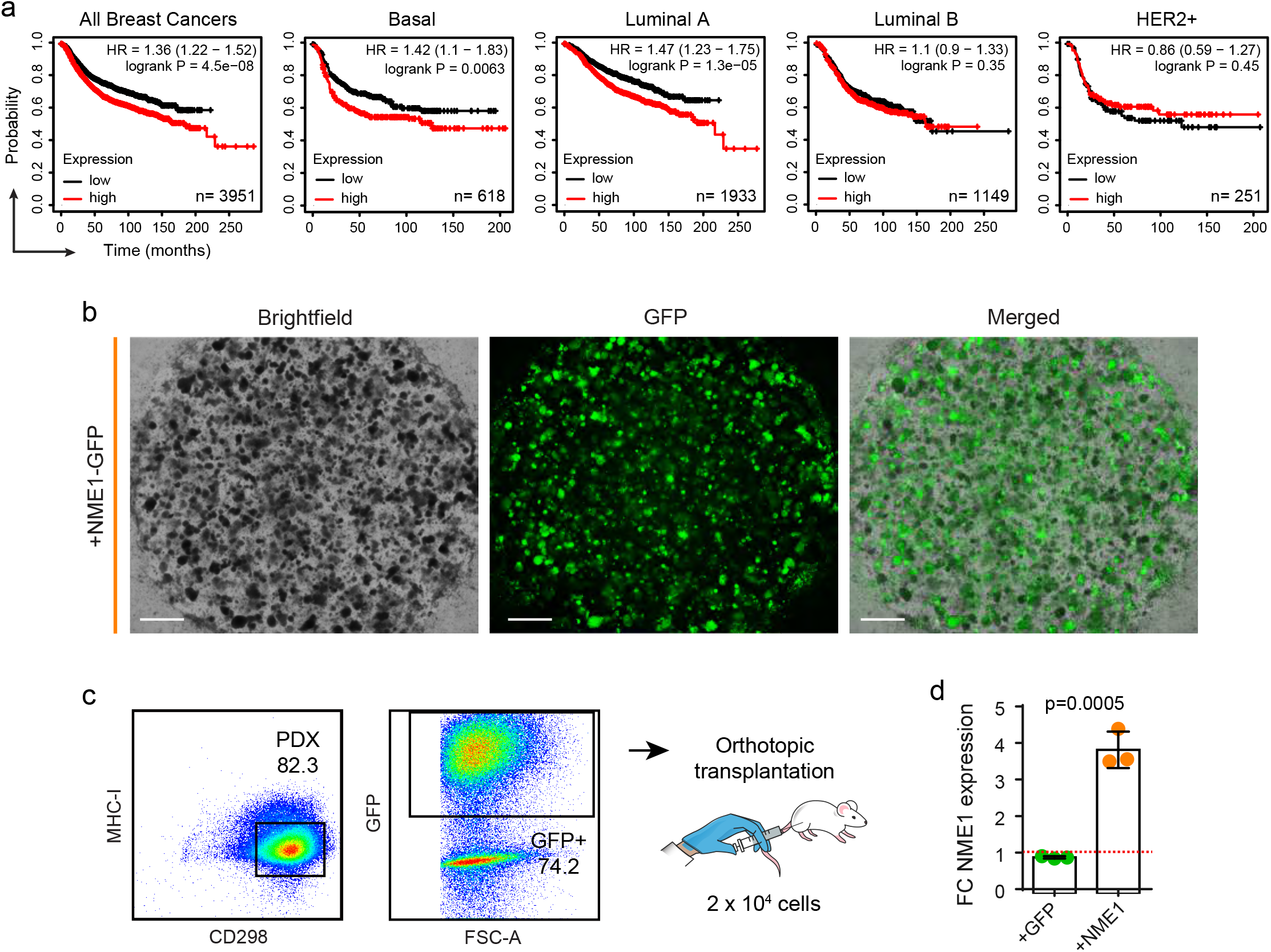
NME1 overexpression in cultured PDX cells and its relevance in patient breast cancer prognosis. **(a)** Kaplan-Meier (KM) curves show relapse free survival (RFS) in breast cancer patients by tumor subtype using the KM plotter database(ref), based on their primary tumor expression of NME1. *P-*values were determined via a log-rank test. **(b)** Images show GFP expression in PDX spheres cultures following lentiviral transduction *in vitro.* HCI010 sphere cells were transduced with +NME-GFP lentivirus at MOI=25 and re-seeded in MAT-E culture at a density of 2 × 10^5^ cells per well. Images show z-stack micrographs of individual wells containing P1 spheres 3 weeks later. Scale bar= 800 µm. **(c)** Flow cytometry plots show gating strategy for sorting transduced human GFP^+^CD298^+^MHCI^−^ PDX cells for orthotopic transplantation and qPCR analysis. **(d)** Bar graph shows qPCR quantification of NME1 expression in control (+GFP) and NME1-GFP (+NME1) transduced cells. Values are plotted as fold change (FC) in NME1 expression in +NME1 relative to +GFP cells and represented as mean ± s.d. *P*-value determined by unpaired t-test.

